# A standard area diagram for Fusarium yellows rating in sugar beet (*Beta vulgaris L.)*

**DOI:** 10.1101/2024.04.23.590831

**Authors:** Olivia E. Todd, Linda E. Hanson, Kevin Dorn

**Affiliations:** USDA-ARS, 1701 Center Ave, Fort Collins, CO, 80525, USA; USDA-ARS, 494 PSSB. MSU, East Lansing, MI, 48824, USDA

**Keywords:** *Fusarium oxysporum*, Fusarium yellows, *Fusarium commune*, Foliar Disease, Disease rating, vascular wilt

## Abstract

Members of the *Fusarium oxysporum* species complex are pathogens of sugar beet causing Fusarium yellows. Fusarium yellows can reduce plant stand, yield, and extractable sugar. Improving host plant resistance against *Fusarium*-induced diseases, like Fusarium yellows, represents an important long-term breeding target in sugar beet breeding programs. Current methods for rating Fusarium yellows disease severity rely on an ordinal scale, which limits precision for intermediate phenotypes. In this study, we aimed to improve the accuracy and precision of rating Fusarium yellows by developing a standard area diagram (SAD). Two SAD versions were created using images of sugar beets infected with *Fusarium oxysporum* strain F19. Each version was tested using inexperienced raters. Comparing both the pilot and improved version showed no statistical differences in Lin’s Concordance Correlation Coefficient (LCC) values to assess accuracy and precision between the two versions (Cb = 0.99 for both versions, ρ*_c_* = 0.97 and 0.96 for version 1 and 2, respectively). In addition, five naïve Bayesian machine learning models which used pixel classification to determine disease score, were tested for congruency to human estimates in version 2. Root mean square error was lowest compared to the “true” values for the unweighted model and a model where necrotic tissue was given a 2x weight (12.4 and 12.6, respectively). The creation of this standard area diagram enables breeding programs to make consistent, accurate disease ratings regardless of personnel’s’ previous experience with Fusarium yellows. Additionally, more iterations of pixel quantification equations may overcome accuracy issues for rating Fusarium yellows.

## 1. INTRODUCTION

Sugar beet (*Beta vulgaris* L.) is one of two economically important sugar crops worldwide. In the United States, 1.1 million acres are raised annually, with the majority of the acreage lying in the great plains and the far west (USDA-ERS, 2021, Commission, 2023). Sugar beet is subject to a number of diseases (Harveson et al., 2009) including Fusarium yellows (also called Fusarium wilt), caused by *Fusarium oxysporum,* and some other diseases caused by *Fusarium* species with similar symptoms (Harveson et al., 2009), that can impact crop and sugar yield. In the United States, losses to *Fusarium*-induced diseases cause $20-90 million dollars in financial loss across 250,000 acres (BSDF, 2021).

The delineation between members of the *Fusarium* species complex has long been up for debate (Skovgaard et al., 2003). While *Fusarium commune* has been identified as a separate sister taxon to *Fusarium oxysporum*, the F19 strain used in this study was thought to belong to *F. commune*, but lacks polyphialides that define the species (unpublished data). Herein we will be referring to the F19 strain as a part of the *Fusarium oxysporum* taxon.

*Fusarium oxysporum* includes many pathogenic and non-pathogen strains that reside in the soil (Gordon and Martyn, 1997). The various formae speciales of *F. oxysporum* cover a broad host range and cause significant crop injury and loss (Edel-Hermann and Lecomte, 2019, Harveson and Rush, 1997). In sugar beet, *F. oxysporum* forma specialis *betae* (FOB) strain F19 causes Fusarium yellows and in susceptible plants can causing wilting, chlorosis of the leaves, stunting, and plant death (Hanson et al., 2009). Because soil-borne fungi are difficult to control with chemical fungicides, a major line of defense against Fusarium yellows in beet is resistant cultivars (El-Aswad et al., 2023). A limited number of sugar beet germplasm with improved resistance to Fusarium yellows has been developed by USDA-ARS pre-breeding programs (Panella et al., 2015, Todd et al., 2023), which have utilized wild germplasm from the USDA National Plant Germplasm System for *Fusarium* resistance trait discovery and introgression. Disease screening is an integral part of the pre-breeding process, highlighting the importance of reliable and repeatable resources to aid in disease screening.

Standard area diagrams (SADs) are designed to visually estimate plant damage, often due to disease (Belan et al., 2020, Araújo et al., 2019, Del Ponte et al., 2017). SADs aid workers of various experiential background in the field when assessing plant damage ranging from whole-plant to single leaf symptoms (Vereijssen et al., 2003). Properly designed SADs have been shown to produce accurate and precise measures of disease when compared to the true disease score (Franceschi et al., 2020, Moreira et al., 2019). Notably, a SAD does not currently exist in sugar beet for *Fusarium*-induced diseases, therefore the aims of this study were to (1) develop a practical, useful and uniform standard area diagram for Fusarium yellows rating on sugar beet using expert raters as “trued” values, and to (2) assess the possibility of applying quick imaging and naïve bayes pixel quantification and imaging as a high throughput, equally accurate method for rating *Fusarium*-induced diseases.

## 2. MATERIALS AND METHODS

### 2.1 Plant material and image acquisition

Plant images were selected from a pre-existing image library of color images (RGB images) of sugar beet plants treated with strain *F. oxysporum* strain F19 (FOB19). All images in the RGB image library were taken of plants treated as follows in controlled, batched experiments. Sugar beet seeds from the accession 20161016 (Fort Collins USDA-ARS unreleased germplasm) that is susceptible to FOB19 were sewn in bleached pots containing steam pasteurized peat-based potting mix (Featherweight Champion, Paonia Soil Co., Paonia, Colorado). The greenhouse was kept at 25C with a photoperiod of 16h. Plants were thinned to one plant per pot after emergence and watered daily. Overhead misters were used to keep relative humidity in the greenhouse at 70%. FOB19, provided by the United States Department of Agriculture-Agriculture Research Service, East Lansing, MI was plated on 100mm plates containing potato dextrose agar (Fisher Scientific, Hampton, NH, USA). Plates were incubated in the dark at 25C for 7 days. Three 5mm^3^ pieces of agar containing actively growing hyphae were aseptically transferred to 150mL of a carboxymethyl cellulose (CMC) liquid culturing medium, shaken at 200 rpm at 25C in the dark for 48-72h (Lai et al., 2020). When plants were at the 4-6 leaf growth stage, multiple 300ml spore suspensions were prepared by filtering the liquid culture through 5-7 layers of sterile cheesecloth. Spore concentration was determined using a hemacytometer and the final suspension was diluted with CMC to 5×10^4^ spores/mL. Plants were treated by soaking roots in the spore suspension for five minutes, while lightly agitating the beaker containing the spore suspension and plants. The negative control was prepared with CMC medium and deionized water to match the spore suspension volume. Plants were pulled from their pots and the roots were gently washed in tap water. Thirty plants were treated per beaker of root dip solution to prevent spore dilution from water held in the roots from washing. After treatment, plants were carefully repotted. As treated plant symptoms progressed, plants were set on a white background and aerially imaged using a 50-megapixel standard smartphone camera every week for four weeks. The image library contains more than 200 images of plants batch-treated with this method.

### 2.2 SAD version development and data collection

The pilot standard area diagram (BvFus-SAD-v1) was an eight-category scale created to represent *Fusarium-*induced disease development based on progression of foliar yellowing, wilting and necrosis and included a written table of disease progression symptoms correlating with a 0-100% scale. Expert breeders and pathologists agreed on the increasing disease severity of the eight images in BvFus-SAD-v1, and manually assigned disease scores on a 0-100% scale. The SAD and the written table were provided to twelve participants who were asked to rate 75 random images from the image library (see data availability statement). The “trued” rating values for BvFus-SAD-v1 were determined by the averaged expert raters’ scores of the 75 images.

In an effort to improve BvFus-SAD-v1 following error analysis and participant feedback on general useability, we followed several guidelines outlined by Del Ponte et al. (2017). BvFus-SAD-v2 was comprised of a ten-category scale created using images from the image library described above to represent disease development with regard to foliar yellowing, necrosis and wilting. We assigned one image per 10% disease interval. The image set was increased to 90 images that more accurately represented the range of symptoms from the image library for participants to rate (see data availability statement). The images were presented in a random order to participants. The “trued” ratings for BvFus-SAD-v2 were determined by the average of three expert ratings. Seventeen participants were asked to assign a disease percentage between 0-100% for each image, and were only provided with a BvFus-SAD-v2 diagram, not an additional table of written symptoms. Participants were given four weeks to complete all 90 ratings.

### 2.3 Naïve Bayes pixel classification with PlantCV

To assess how well using machine learning models were able to assign disease scores compared to the human ratings in BvFus-SAD-v2, naïve bayes pixel level plant segmentation with PlantCV (Abbasi and Fahlgren, 2016) was used to quantify diseased tissue. Five plant images from the image library were selected for pixel sampling, and more than 300 RGB values representing “Healthy”, “Chlorotic”, “Necrotic”, and “Background” categories were manually assigned using ImageJ (https://imagej.net/ij/index.html) and input into an excel file under the appropriate classifier (see data availability statement). Using the PlantCV method, all 90 images used in BvFus-SAD-v2 were given a quantitative disease percentage score based on the total pixels from the “Necrotic” and “Chlorotic” categories following the equation or a derivation of the equation below:

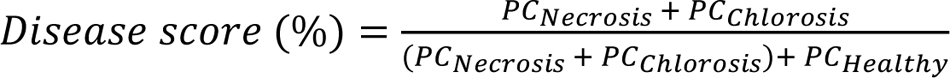

Where *PC* = Pixel count for the subscript Bayesian classifiers “Chlorosis”, “Necrosis” and “Healthy”.

Several machine learning models were iterated over the 90 images in the BvFus-SAD-v2 image set, “trued” to the three expert raters’ disease scores. Four weighted adjustments to the equation described above were made, where the *PC*_*Necrosis*_ value was given a 2x multiplier, the *PC*_*Chlorosis*_ value was given a 2x multiplier and a 5x multiplier in separate models, and in a fourth model where *PC*_*Chlorosis*_ and *PC*_*Necrosis*_ values were both given 2x multipliers. The fifth model was unweighted, displayed as the above equation.

### 2.4 Data processing and statistical analysis

Each image selected for BvFus-SAD-v1 and BvFus-SAD-v2 of the standard area diagram received a “trued rating” using the mean of three expert raters’ assigned disease scores.

Participant rated values were “estimated” values. Data analysis was conducted using R version 4.2.0 and modified scripts from Franceschi et al. (2020). A generalized linear mixed model was used to estimate the mean of each Lin’s concordance correlation (LCC) parameter for comparison of BvFus-SAD-v1 and BvFus-SAD-v2. The R package “lsmeans” was used to compare the mean disease score estimates based on Tukey’s HSD test. The agreement between rater estimates in relation to the “true” values in BvFus-SAD-v1 and BvFus-SAD-v2 was calculated using statistical parameters *r*, a measure of the degree of scatter around a line of best fit and *C_b_*, the bias correction factor, from LCC. Based on these two parameters, the concordance correlation coefficient ρ_*c*_ is defined as

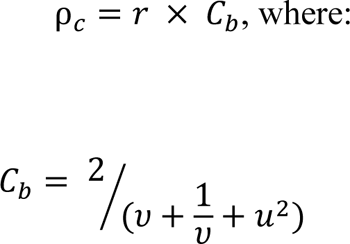

And

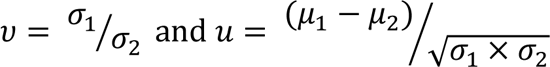

The means for both “estimated” and “true” values are represented by μ_1_and μ_2_, respectively. The standard deviations for the “estimated” and “true” values are represented by σ_1_and σ_2_, respectively. The term ν measures the scale shift between “estimated” and “true” values, and *u* represents location shift relative to the scale. The ρ_*c*_ term measures the variation of data from a 45-degree line (with a slope of 1 and intercept of 0) (Lawrence and Lin, 1989, Nita et al., 2003). A table of values for the above terms are provided for each SAD version (Table 2). Data were graphed using R and GraphPad Prism version 10.0.2 for MacOS (GraphPad Software, Boston, MA, USA, www.graphpad.com).

To analyze the difference between the five machine learning models and the human estimates in BvFus-SAD-v2 (model names C2, C5, N2, N2C2, No Weight and Human), 95% confidence intervals and Root Mean Squared Error were generated for each machine learning model as well as the human estimates using Prism and R respectively.

## 3. Results

### 3.1 Statistical comparison of accuracy of the two BvFus-SAD versions

BvFus-SAD-v1 contained 8 levels of *Fusarium-*induced foliar disease, with the 0% value representing no disease and 100% representing complete death of the root and necrosis of the leaves (Figure 1). Participants were also provided with written descriptions of ten symptom categories to supplement the 8 images used to make BvFus-SAD-v1 (Table 1). BvFus-SAD-v2 improved upon BvFus-SAD-v1 by adding an image at three additional disease percentage categories at 20%, 40% and 60%, giving it eleven total images (Figure 2A), and removed the written table for ease of use. Participants were not provided the naïve bayes false colorized images with machine learning assigned disease ratings estimates as determined by the unweighted equation in section 2.2 (Figure 2B).

**Figure 1:**
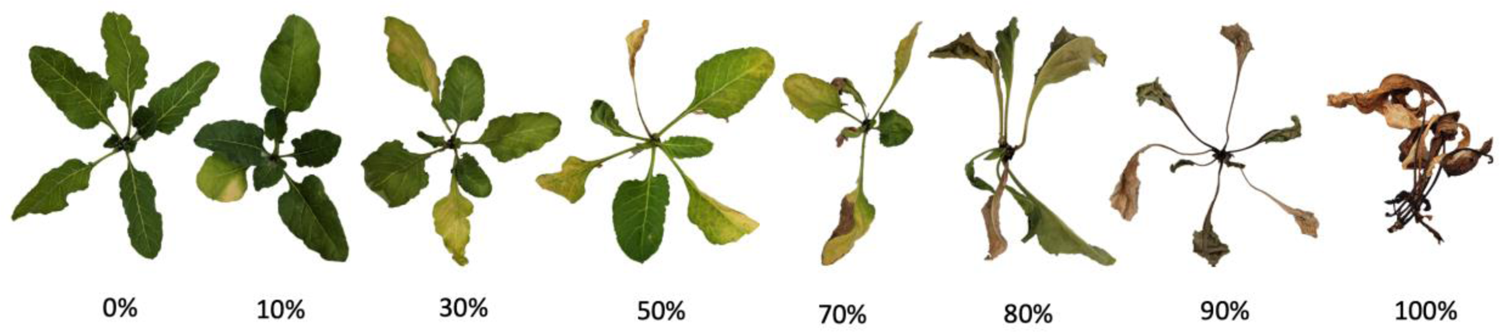
Standard area diagram for progression of Fusarium yellows foliar injury in sugar beet, BvFus-SAD-v1. Eight categories were assigned by expert raters.

**Figure 2:**
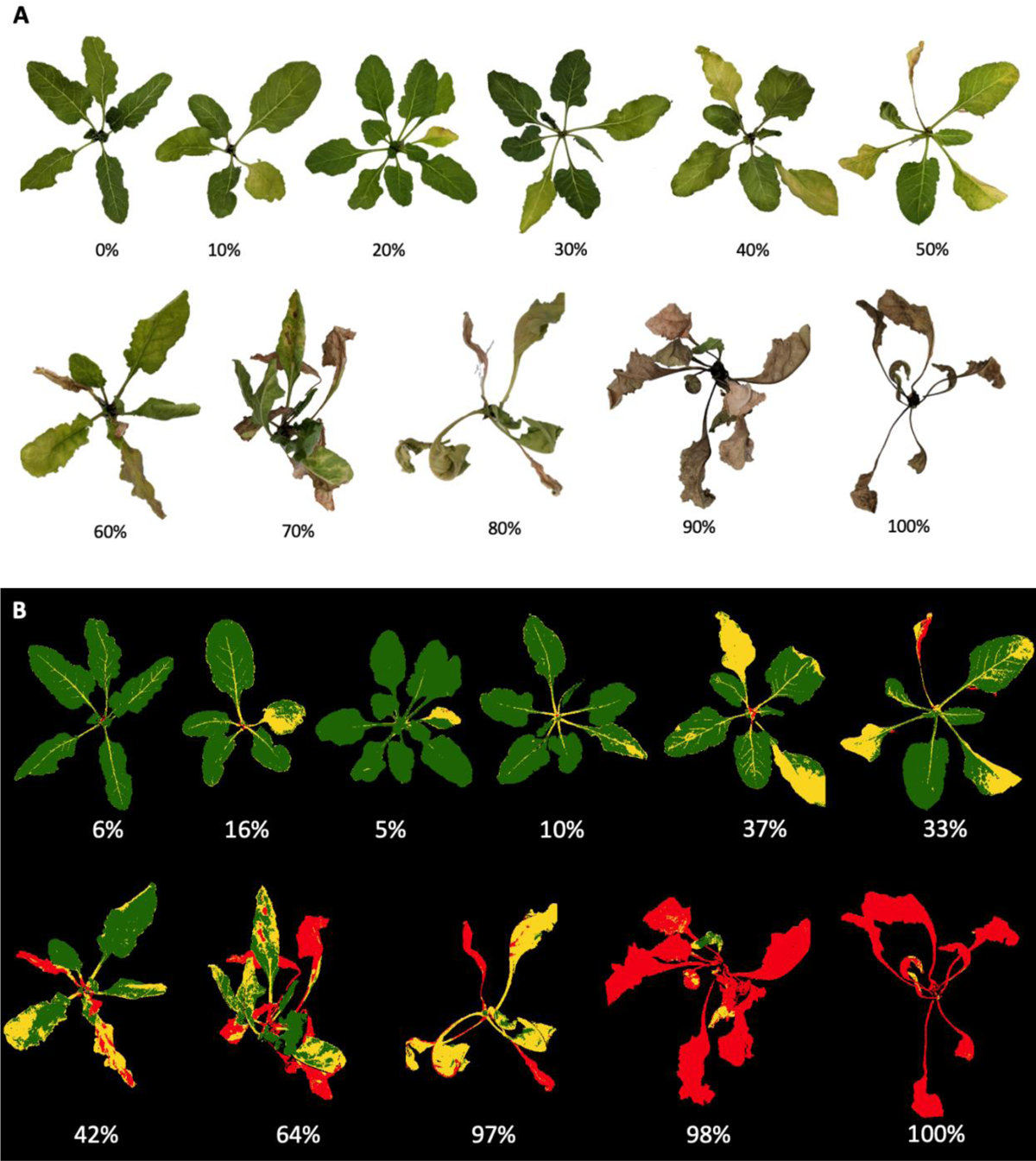
A. BvFus-SAD-v2 of the Fusarium standard area diagram for Fusarium yellows foliar injury in sugar beet. B. Machine learning classified injury scales (as a percent) based on RGB imaging and pixel quantification into Bayesian classifiers. Color scale: green = healthy tissue, yellow = chlorotic tissue, red = necrotic tissue.

**Table 1.** Categorical plant injury descriptions for Fusarium yellows of sugar beet provided alongside the pictorial BvFus-SAD-v1 diagram (Figure 1A).

| Rating Category | Category Description |
| --- | --- |
| 0 | All leaves are green and show no signs of yellowing, necrosis, or wilting |
| 10 | 1-2 older leaves show signs of yellowing, new leaves are green, no wilting |
| 30 | 2-3 older leaves yellowing, new leaves are green, wilting may be seen |
| 40 | 2-3 older leaves yellowing, new leaves may have some yellowing, wilting may be seen, no necrosis |
| 50 | Older leaves may show advanced signs of yellowing, wilting or slight necrosis. New leaves may begin to show signs of yellowing or wilting. No necrosis of new leaves |
| 60 | Older leaves show advanced signs of yellowing, wilting or necrosis. New leaves may begin to show signs of yellowing, wilting or necrosis |
| 70 | Older leaves may be fully necrotic. New leaves show signs of yellowing, wilting or necrosis |
| 80 | Both older and newer leaves may show signs of advanced wilting and necrosis. At this stage, there is little to no yellowing |
| 90 | Both older and newer leaves are fully necrotic, but still attached to the base of the crown |
| 100 | All leaves are fully necrotic and may be detached from the crown which may be undetectable or not present. White fungal hyphae may be visible at the base of the leaves |

Similar to BvFus-SAD-v1, 0 and 100% represented no disease and complete death of the plant, respectively. Despite these improvements, all tested parameters of LCC (*C_b_*, ρ_*c*_, ν, *r*, and *u* were statistically unchanged, with the exception of scale and location shift (ν and *u*), which were not different enough to have an overall effect on the correlation coefficient (*C_b_*) (Table 2). To further improve BvFus-SAD-v2, we chose a representative set of images for participants to rate which represented the full range of symptoms instead of randomly selecting the images from the image library. This enriched the spread of the estimated severity distribution significantly from BvFus-SAD-v1 to BvFus-SAD-v2, filling necessary gaps around 20, 40 and 60% (Figure 3A) and allowed us to accurately assess any areas of inconsistency in rating across the entire scale.

**Table 2:** Lin’s concordance correlation coefficient parameters compared by lsmeans in R between two standard area diagrams for Fusarium yellows of sugar beet, BvFus-SAD-v1 and BvFus-SAD-v2. All comparisons by column were compared by a Tukey HSD test, indicating significant differences in scale and location shift, components of the overall correlation coefficient (*C_b_*).

| | Scale shift ($v$) | Location shift ($u$) | $C_b$ | $r$ | $\rho_c$ |
| --- | --- | --- | --- | --- | --- |
| BvFus-SAD-v1 | 0.95 ($\pm 0.02$)<sup>a</sup> | 0.05 ($\pm 0.02$)<sup>a</sup> | 0.99 ($\pm 0.002$)<sup>a</sup> | 0.97 ($\pm 0.007$)<sup>a</sup> | 0.97 ($\pm 0.004$)<sup>a</sup> |
| BvFus-SAD-v2 | 1.00 ($\pm 0.01$)<sup>b</sup> | -0.01 ($\pm 0.02$)<sup>b</sup> | 0.99 ($\pm 0.002$)<sup>a</sup> | 0.96 ($\pm 0.005$)<sup>a</sup> | 0.96 ($\pm 0.005$)<sup>a</sup> |

**Figure 3:**
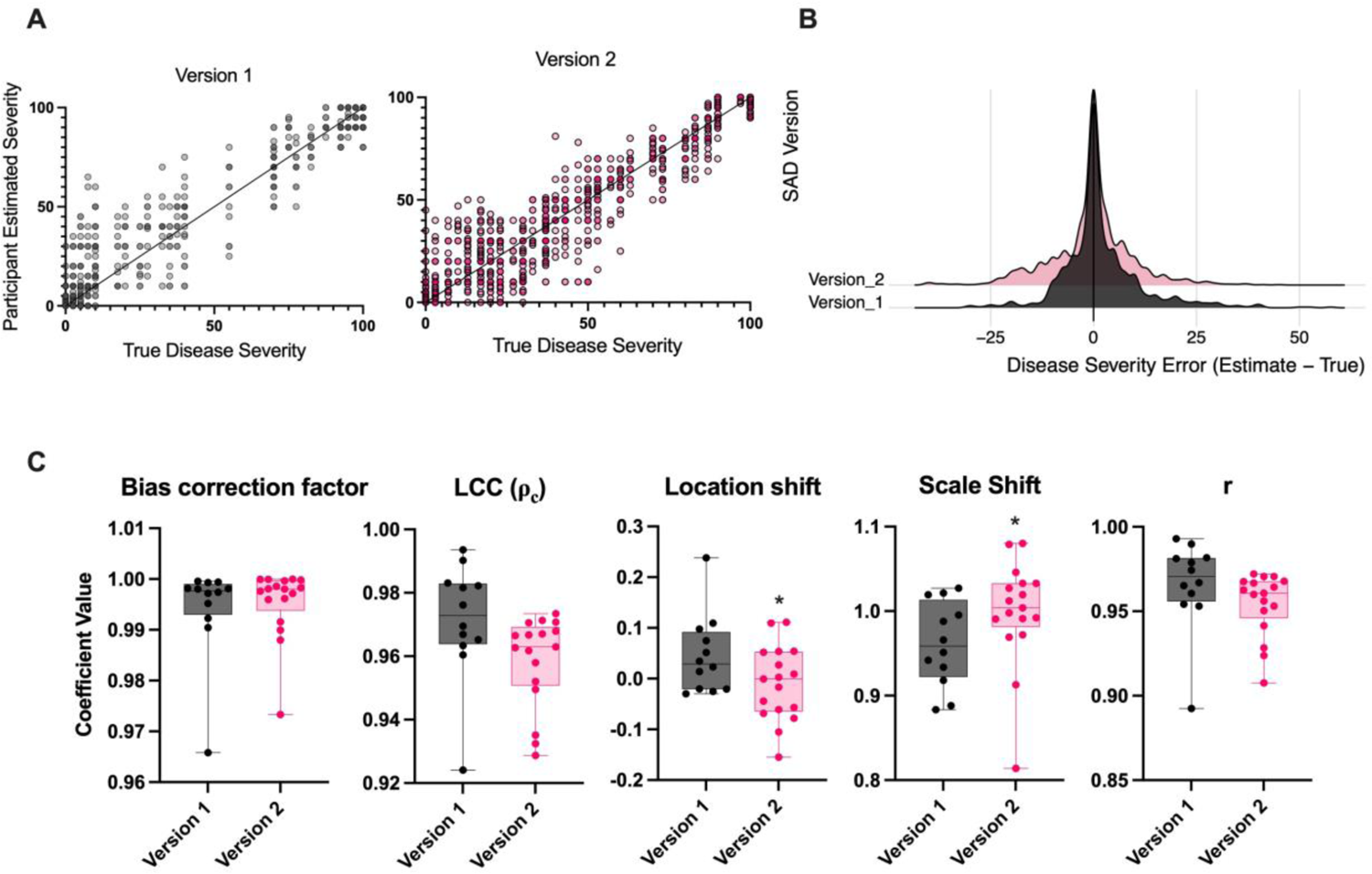
A. Participant estimated severity versus true severity. True severity was determined by the mean of two expert raters for BvFus-SAD-v1 (left panel) and three expert raters for BvFus-SAD-v2 (right panel). The solid black line indicates a point in which the rater estimate matches the true value perfectly (Slope = 1, intercept = 0). B. Disease severity error term (estimated value minus true value) for both versions of the standard area diagram. C. Lin’s concordance correlation coefficient (LCC) comparisons for BvFus-SAD-v1 and BvFus-SAD-v2. Statistical significance between LCC parameters are represented for * p<0.05 Tukey HSD comparisons.

Both BvFus-SAD-v1 and BvFus-SAD-v2 had a relatively normally distributed disease severity error curve (Figure 3B). Both SADs had high *r* and ρ_*c*_ ranges above .90 for all individual participants, indicating a high level of accuracy and reproducibility (Figure 3C). Participants using BvFus-SAD-v1 showed more consistent overestimation (by 5-15%) on images that had a true disease score between 0-75%% in BvFus-SAD-v1, where BvFus-SAD-v2 participants overestimated true disease values between 0-25%, and underestimated values from 25-40% (SI Figure 1).

### 3.2 Comparing machine learning ratings to BvFus-SAD-v2 human estimates

We compared parameters of the five machine learning models “trued” to the expert raters’ values and the human estimates of BvFus-SAD-v2. We endeavored to find a model that matched or outperformed the human estimates, indicating its possible use for future automation studies. While there was only one estimate per image for each of the machine learning models, there were multiple images that had the same “true” rating value. Each model showed a widespread dispersion of the error term; the majority of the models overestimated disease severity (Figure 4B). Raw disease severity estimates show that the unweighted model (NoWeight) and the model with the “necrosis” classifier weighted 2x (N2), minimized the error term compared to the “true” rating values (Figure 4C). All five of the models overestimated the disease scores from 0-20% highlighting a consistent error in the modeling system (Figure 4D). The NoWeight and N2 models mimic the human estimates between 25-50% disease severity, where disease severity is consistently underestimated compared to the “true” disease rating (Figure 4D). The confidence interval and linear line of best fit for the NoWeight and N2 models most closely resembled the trend of the human estimates (Figure 5). Root Mean Squared Error (RMSE) was smallest for the Human estimates (9.85) indicating the closest fit for the estimate regression line to the “true” values. The NoWeight model and the N2 model both had RMSEs of ∼12.5 (Table 3).

**Figure 4.**
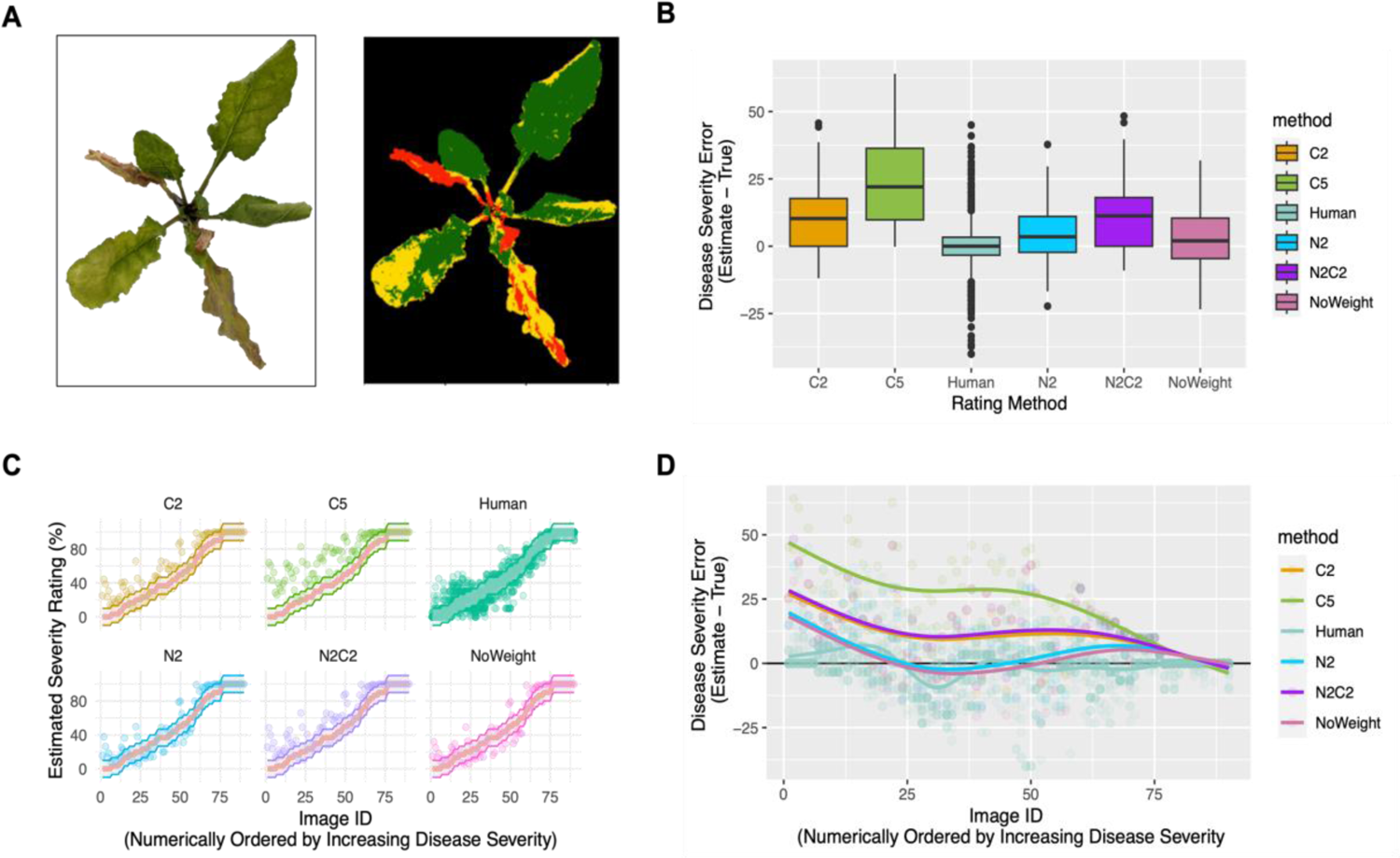
A. Sugar beet plant infected with *Fusarium oxysporum* strain F19, symptomology (right panel) and false colorized image (left panel) based on disease categories “healthy” in green, “chlorosis” in yellow, “necrosis” in red and “background” in black. B. Boxplots displaying the spread of the disease severity error for all six disease rating methods (five weighted machine learning models and the BvFus-SAD-v2 human estimates). C. Estimated disease severity per rating method with a 95% CI. The red line represents the true disease ratings as determined by the mean of three expert raters for each of the 90 plant images. D. Disease severity error for each of the 90 plant images per disease rating method.

**Figure 5:**
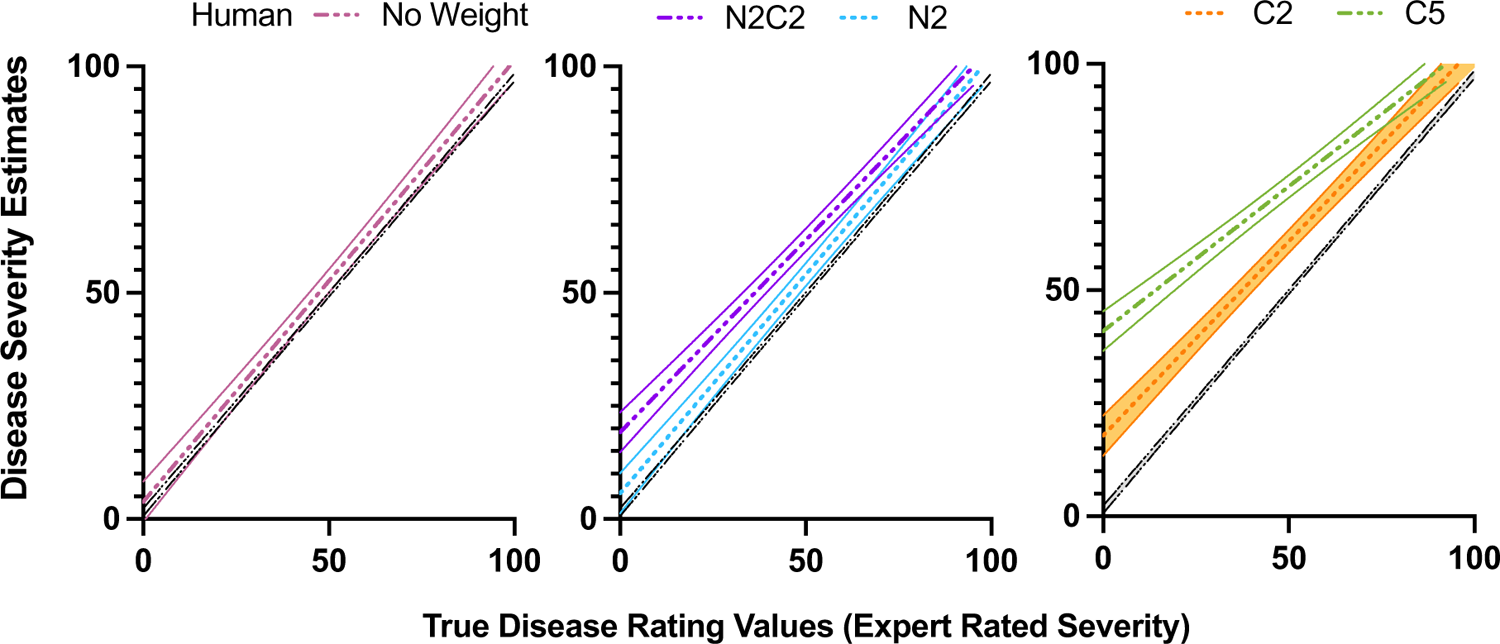
95% confidence intervals for disease severity estimates versus expert rated disease values per machine learning model and the BvFus-SAD-v2 human estimates. Human estimates are provided in each subsection comparison.

**Table 3:** Root Mean Square Error (RMSE) and R^2^ value for all 5 machine learning models and human estimates in the BvFus-SAD-v2 dataset for Fusarium yellows of sugar beet.

| Model | RMSE | R squared value of the fit line |
| --- | --- | --- |
| Human | 9.85 | 0.92 |
| NoWeight | 12.40 | 0.88 |
| N2 | 12.65 | 0.88 |
| C2 | 16.86 | 0.86 |
| C5 | 28.74 | 0.78 |
| N2C2 | 17.46 | 0.86 |

## Discussion

Generally, both BvFus-SAD-v1 and BvFus-SAD-v2 are reliable and easily reproducible by participants with no prior experience with rating Fusarium yellows in sugar beet. Because there were no statistically significant improvements on the LCC parameters between BvFus-SAD-v1 and BvFus-SAD-v2, either version can be used based on the specific use cases or unique needs. However, because BvFus-SAD-v2 used an expanded the image library, has categorical images at 10% intervals, we recommend the use of BvFus-SAD-v2 for Fusarium yellows screening (grayscale version SI Figure 2). While we did not test the effect of pre-training human raters to gain experience in Fusarium yellows symptomology before assessing disease score, there is evidence to suggest that familiarity with the specific disease symptomology does significantly improve accuracy and precision of ratings (Franceschi et al., 2020, Bock et al., 2009, Yadav et al., 2013). Additionally, it should be understood that due to the overestimation generally caused by BvFus-SAD-v1, use of the BvFus-SAD-v1 in breeding programs may cause accidental culling of resistant germplasm. Conversely, the underestimation in BvFus-SAD-v2 specifically in the 25-40% range was by approximately 10% (one category scale). Because these rating trends are more conservative between the 25-40% categories, this may cause accidental selection of weakly resistant germplasm. The head user of the diagram may consider the disease severity cutoff for their program when selecting which SAD to use.

After evaluating the ease of use for training the machine learning models with RGB image classification, we assess that there is potential to improve the use of naïve bayes classifiers with a machine learning pipeline for Fusarium yellows disease rating. While using pixel quantification classifiers has been shown to be successfully implemented for disease quantification on individual leaves (Mondal et al., 2017, Bannihatti et al., 2022), whole plant imaging studies in dicots are lacking (reviewed by (Humplik et al., 2015). Additionally, some of the symptoms of Fusarium yellows disease progression are difficult to categorize via RGB pixel color (such as wilt), which is a major limitation of using a pixel classification method and is an important symptom with this disease (Ruppel, 1991). Utilizing multifaceted, advanced imaging systems such as chlorophyll, multispectral, and hyperspectral imaging allow for a more comprehensive phenotyping strategy, but may be more costly and complicated to implement.

Despite this, these methods have been successful or show promise for identifying disease and classifying disease severity for Cercospora leaf spot, powdery mildew and other beet disease (Chaerle et al., 2007, Mahlein et al., 2012, Adem et al., 2022, Leucker et al., 2016, Barreto et al., 2020), but these technologies have yet to be optimized for *Fusarium*-induced diseases in sugar beet.

Providing an updated, standard graphic reference for Fusarium yellows in sugar beet is integral for maintaining high quality breeding stock for seed companies and consumers. When using either SAD for Fusarium yellows assessment, the head breeder or researcher should describe to all individuals rating an experiment where the disease severity cutoff is for the research question of interest, especially when considering other potential *Fusarium* management tools. Typically, after 30 days, empirical evidence suggests that resistant plants that fall into BvFus-SAD-v2 <60% disease severity will recover from *Fusarium oxysporum* F-19 infection in the greenhouse when maintained at 28C (Todd et al., 2023, Webb et al., 2015). When conducting high-throughput screening in the greenhouse for Fusarium yellows resistance, pixel quantification using PlantCV may provide a cheap and easy image solution that is most comparable to human estimates from 20-100% disease severity on the BsFus-SAD-v2 scale. For the most accurate results, we recommend human validation for images rated between 0-20%.

## Acknowledgements

The authors would like to acknowledge the raters’ participation in this study. Funding provided by USDA-ARS CRIS projects 3012-21220-011-000-D and 5050-21220-017-000-D.

## Data availability statement

Scripts, images and excel sheets are available on the following Github page: https://github.com/oetodd/Fusarium_standard_area_diagram_2024

## Author contributions

OET Investigation, writing original draft, review and editing; methodology; LH resources, review and editing; KMD review and editing.

## Supporting information legends

Supplemental Figure 1: Disease severity error estimates ordered by increasing true disease severity shows participant overestimation in BvFus-SAD-v1 (Version_1) and overestimation between true disease values 0-25% and underestimation between 25-40% in BsFus-SAD-v2 (Version_2).

Supplemental Figure 2: Greyscale, printable standard area diagram of BvFus-SAD-v2 for Fusarium yellows foliar injury in sugar beet.

