## Supplemental Information for "A standard area diagram for Fusarium yellows rating in sugar beet (*Beta vulgaris L.)*"

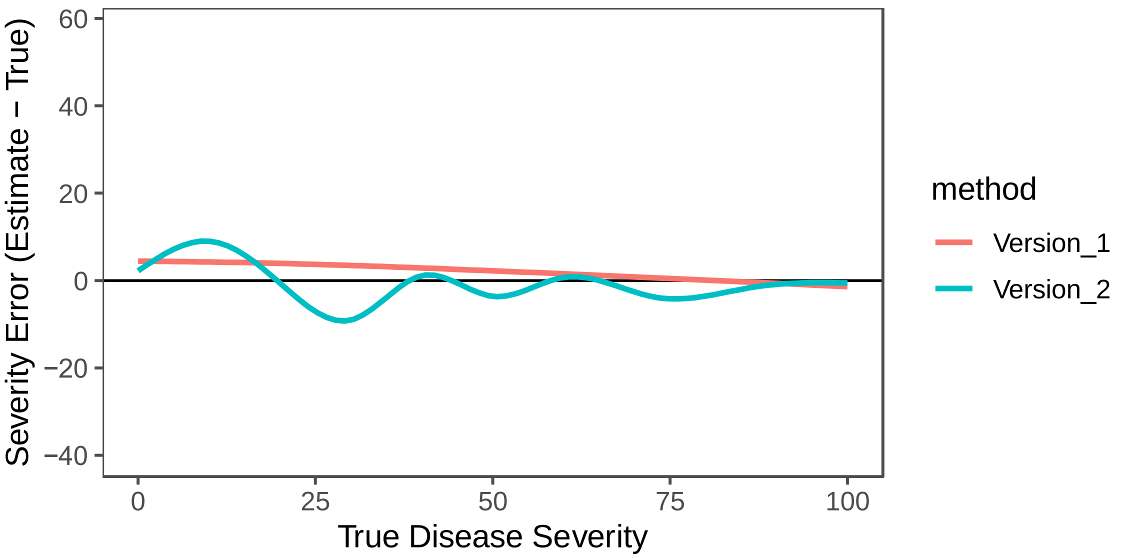


Supplemental Figure 1: Disease severity error estimates ordered by increasing true disease severity shows participant overestimation in BvFus-SAD-v1 (Version_1) and overestimation between true disease values 0-25% and underestimation between 25-40% in BsFus-SAD-v2 (Version_2).

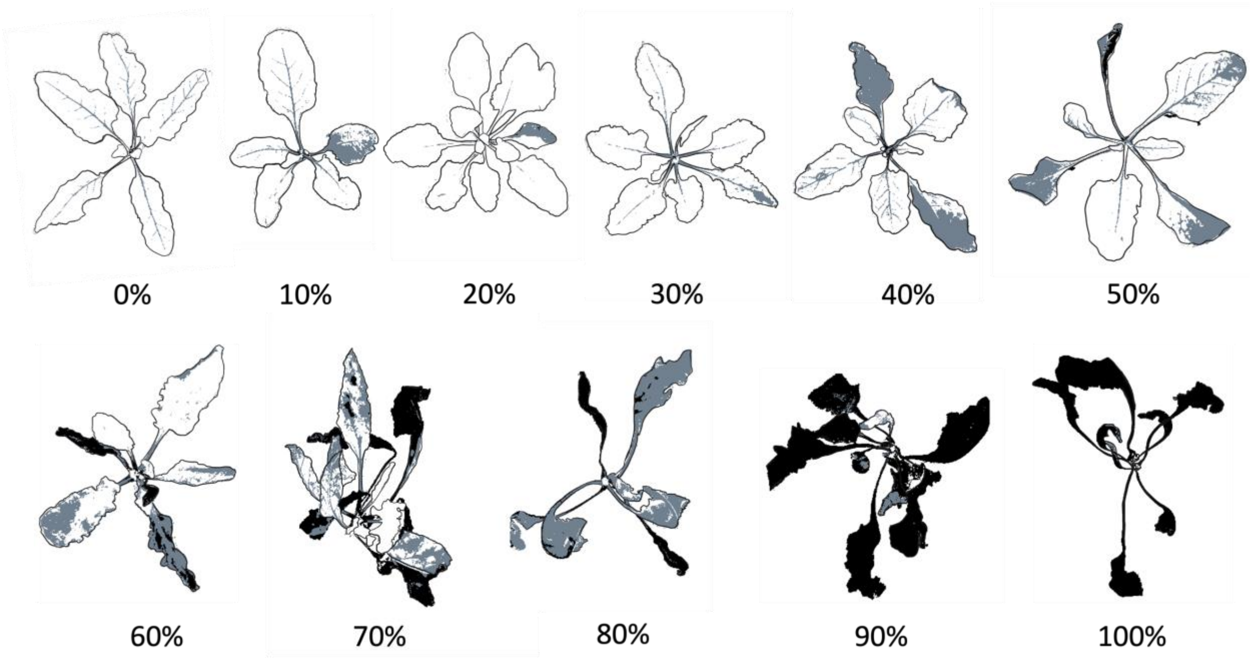


Supplemental Figure 2: Greyscale, printable standard area diagram of BvFus-SAD-v2 for Fusarium yellows foliar injury in sugar beet.
